# The rise of sparser single-cell RNAseq datasets; consequences and opportunities

**DOI:** 10.1101/2022.05.20.492823

**Authors:** Gerard A. Bouland, Ahmed Mahfouz, Marcel J.T. Reinders

**Affiliations:** Delft Bioinformatics Lab, Delft University of Technology, Delft, The Netherlands; Department of Human Genetics, Leiden University Medical Center, Leiden 2333ZC, The Netherlands; Leiden Computational Biology Center, Leiden University Medical Center, Leiden 2333ZC, The Netherlands

## Abstract

There is an exponential increase in the number of cells measured in single-cell RNA sequencing (scRNAseq) datasets. Concurrently, scRNA-seq datasets become increasingly sparser as more zero counts are measured for many genes. We discuss that with increasing sparsity the binarized representation of gene expression becomes as informative as count-based expression. We show that downstream analyses based on binarized gene expressions give similar results to analyses based on count-based expressions. Moreover, a binarized representation scales to 17-fold more cells that can be analyzed using the same amount of computational resources. Based on these observations, we recommend the development of specialized tools for bit-aware implementations for downstream analyses tasks, creating opportunities to get a more fine-grained resolution of biological heterogeneity.

## Introduction

Since its introduction, single-cell RNA sequencing (scRNAseq) has been vital in investigating biological questions that were previously impossible to answer(1–4). Continuous technological innovations are resulting in a consistent increase in the number of cells being measured in a single experiment. However, at the same time, datasets have become sparser, i.e. no counts measured for an increasing number of genes. This sparsity has generally been seen as a problem, especially since standard count distribution models seem to fail in explaining the excess of zeros(5–8). This zero-inflation has sparked discussions about whether the excess of zeros can be explained by mainly technological or biological factors(5, 8–10). As the field moves towards sparser datasets, it is vital to know what the consequences are of the ever-increasing abundance of zero measurements. In 2020, Qui et al. (11) proposed to embrace zeros as useful signal and developed a clustering algorithm requiring only binarized scRNAseq data (a zero representing a zero count and a one a non-zero count). Although this was the first paper explicitly stating zeros (“dropouts”) to be informative, binarization of scRNAseq was already used in practice in 2015 to infer gene regulatory networks(12). Since then, several methods have employed binarized scRNA-seq data. For instance, for dimensionality reduction that improved cell type classification and trajectory inference compared to using counts(13), as well as for differential expression analysis(14).

Using 52 datasets, published between 2015 and 2021, we show that as the sparsity of scRNAseq data increases, the detection pattern of expression (binarized scRNAseq data) becomes as informative as the quantification of expression (counts). We explain this by showing that the driving process behind zero-inflation is mainly explained by biological heterogeneity. Motivated by these findings, we demonstrate that downstream analyses based on solely the detection pattern are in line with those based on quantification. Together with further advantages of binarized scRNAseq data, this opens possibilities in analyzing extremely large datasets in an efficient manner.

### More cells, more zeros

Across the 52 datasets the association between the year of publication and the number of cells shows a Pearson’s correlation coefficient of r = 0.57 (**Fig. 1a**). For instance, the average dataset in 2015 (n = 7) had 704 cells while the average dataset in 2020 (n = 5) had 68,100 cells. One consequence of having more cells is that datasets are becoming sparser, exemplified by a Pearson’s correlation coefficient of r = -0.72 (**Fig. 1b**) for the association between the number of cells and detection rate. It is likely that this trend will continue over the next years as having a larger number of cells has statistical benefits(15, 16).

**FIGURE 1:**
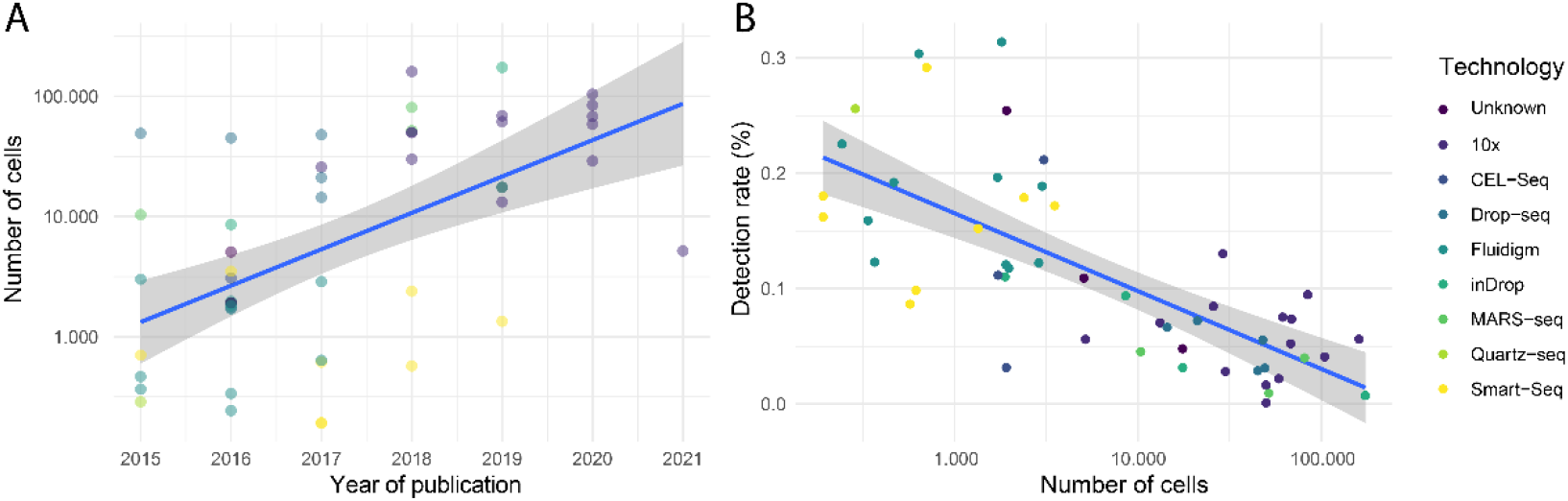
Using R-package scRNAseq (v2.8.0), 52 datasets ranging in date of publication from 2015 to 2021 were downloaded. (a) Scatterplot of the number of cells (log scale) against the date of publication. (b) Scatterplot of the detection rate (%) (y-axis) against the number of cells (log scale, x-axis).

### More zeros, less signal in expression counts

As zeros become more abundant, the detection pattern might be as informative as the expression counts. Using cells from four datasets, with varying degrees of sparsity, we observed a strong correlation (Pearson’s ***r*** ≥ 0.73) between the log-normalized expression counts of a cell and its respective binarized variant (**Fig. 2**). This strong correlation implies that the binarized signal captures most of the signal present in the log normalized count data. Interestingly, there was a strong correlation between this correlation (between counts and binarized profiles) and the detection rate (**Fig. 2**, mean correlation = -0.76). This indicates that as datasets become sparser, the quantification of expression becomes less informative with respect to the detection pattern.

**FIGURE 2:**
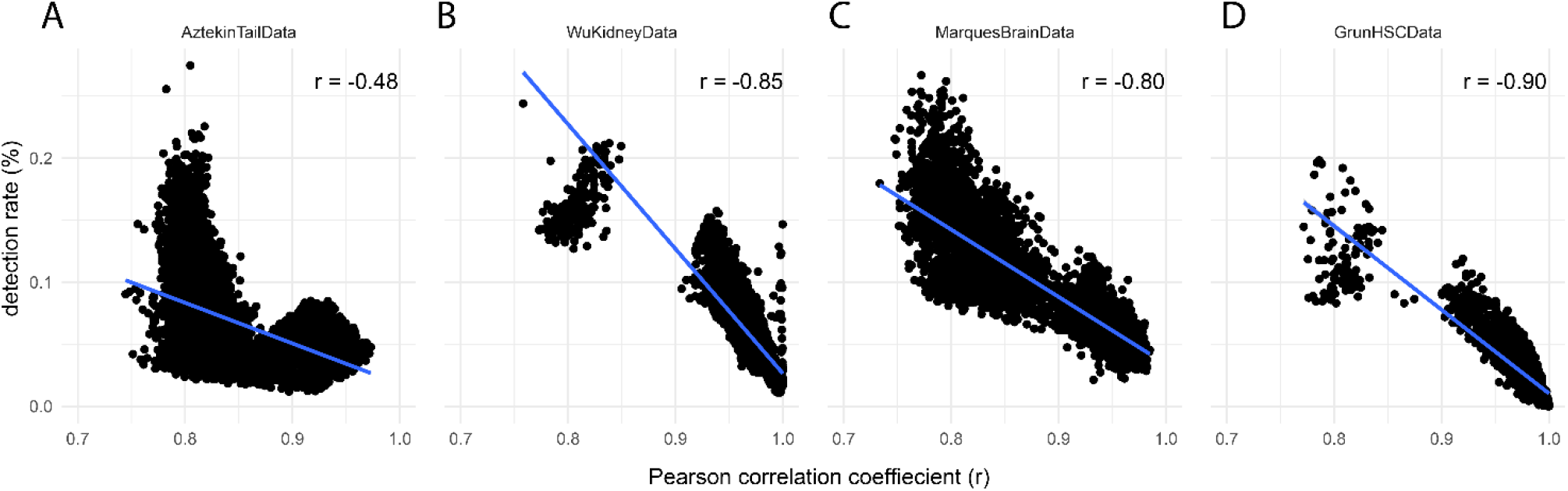
Pearson correlations per cell between log normalised expression profile and the binarized profile across four different datasets (between brackets: detection rate% : median, Q1, Q3) : **a)** AztekinTailData(17) (6.3% 4.6%, 8.8%) **b)** WuKidneyData(18) (4.2%, 3.1%, 5.8%) **c)** MarquesBrainData(19) (10.4% 7.8%, 13.6%) and **d)** GrunHSCData(20) (2.2%, 0.5%, 4.7%). X-axis represents the correlation, Y-axis the (detection rate %) in a cell.

### Expression counts add little to no information on top of the detection pattern

To test whether counts can actually be discarded in practice, we compared the performance of automatic cell-type identification and differential expression analysis methods using counts and binarized data. First, we used two existing automatic cell-type identification methods, scPred and SingleR(21, 22). The median F1-score as well as the global accuracy were very similar between cell-type identifications based on the binarized data and identifications based on the log-normalized count data (**Fig. 3a, Fig. 3b**). This finding implies that the quantification of expression does not add information for cell-type identification. This conclusion was further supported by randomly shuffling the non-zero counts, which resulted in an almost similar cell-type identification performance (**Fig. 3c, Fig. 3d**).

**FIGURE 3:**
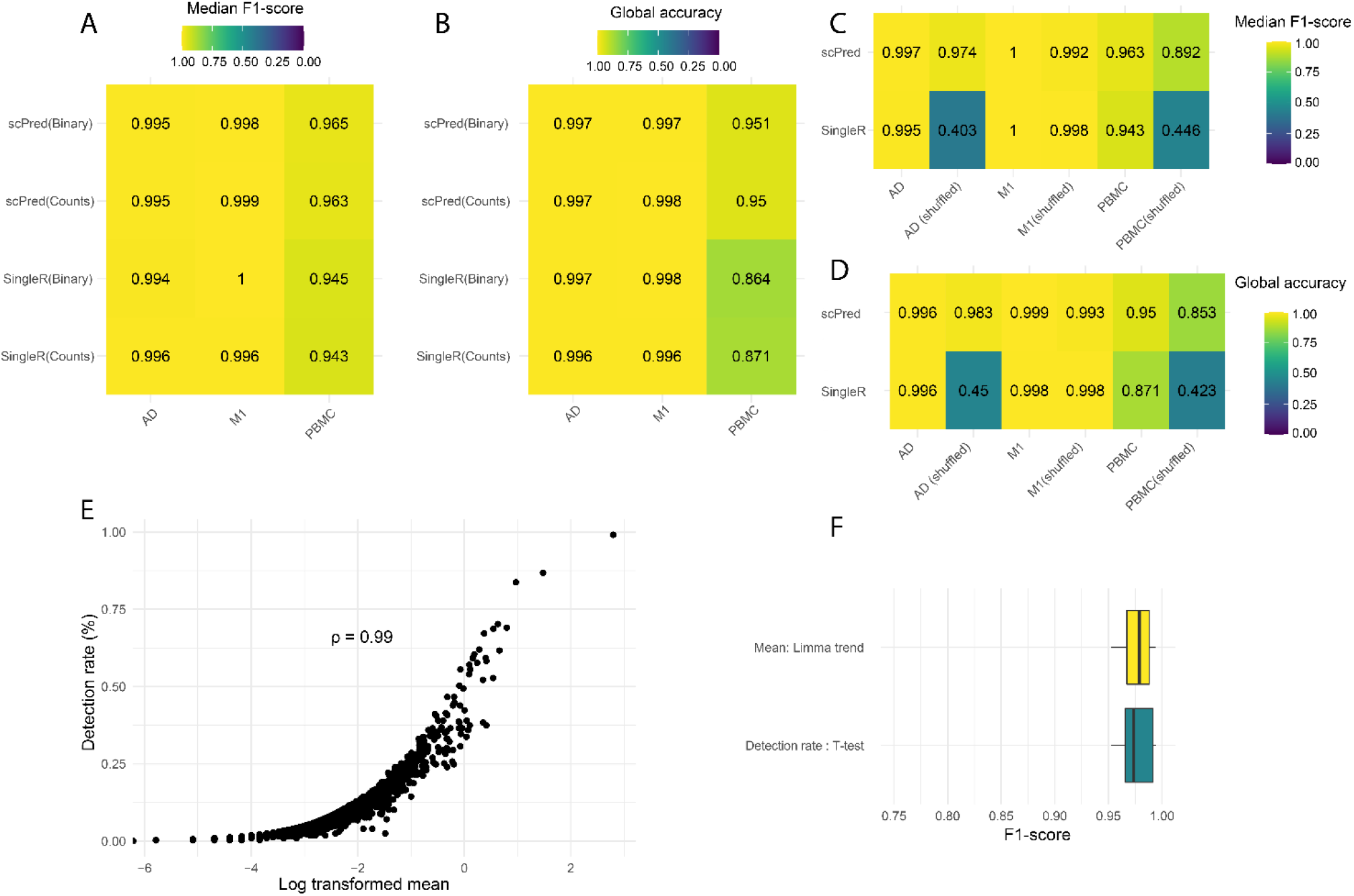
(a,b) The performance of cell type identification by SingleR(21) and scPred(22) when applied to binarized data (Binary) or log-normalized expression (Counts) for three relatively recent datasets: AD(25), M1(26), PBMC(27). (a) show the median F1-score, and (b) the global accuracy score. (c,d) Similar as in (a,b) but now with the non-zero expressions being shuffled. (e) Scatterplot of detection rate (y-axis) vs mean expression(x-axis) for all genes (n = 30.062) of one individual. (f) F1-score indicating how well simulated differentially expressed genes in pseudobulk data can be found back when either count data is used or binarized data.

Next, we evaluated whether counts can also be discarded when pseudobulk data is considered. For each individual, in a dataset containing scRNAseq data of the prefrontal cortex of 34 individuals(23), we generated pseudobulk data by either taking, for each gene, the mean expression across all cells, or the rate of non-zero values across all cells (detection rate). The Spearman’s rank correlation (across all genes) was ≥ 0.99 (**Fig. 3e**) for every individual, indicating that for pseudobulk aggregation, implying that the binarized representation faithfully represents counts. ≥≥To quantify this further, we simulated 10 datasets with muscat(15), each with 50,000 cells and 2,000 genes≥ of which 25% were differentially expressed between two groups comprised each of 10 individuals (2,500 cells per individual). Pseudobulk data for each individual was generated. Using Limma trend(24) for mean gene expression and t-test for the detection rate, we identified differential expressed genes and evaluated whether we found the simulated ones back. The F1-score for the count and binarized data representations were similar(**Fig. 3f**), again indicating that the quantification of expression did not add additional information to test for differential expressed genes in pseudobulk data.

### Zero-inflation is driven by biological heterogeneity

Whether zero-inflation associates with technical or biological origins is heavily debated(8). One compelling reason for this debate is the fact that within a single dataset some genes are zero-inflated, while others are not(5, 8). We argue that this observation is related to whether a gene is only expressed in a subpopulation of cells (e.g. marker genes) or whether a gene has a stable expression (e.g. housekeeping genes). To substantiate our claim, we used BDA(14) to identify the top 100 most differentially expressed genes between two cell populations and the top 100 most stable expressed genes in a 10X dataset(26) as well as a Smart-Seq dataset(28). Next, we applied scRATE(5) to identify the best distribution model for the observed expression of the identified genes, being either a Poisson, a Negative Binomial or their zero-inflated counterparts. A fisher exact test showed that a zero-inflated model was enriched in the top 100 differentially expressed genes, and a non-zero inflated model was enriched in the top 100 stable expressed genes (Table 1). Like earlier work (5), we conclude biological heterogeneity to be the main driver of zero-inflation.

**TABLE 1:** Enrichment of zero-inflated distributions for the top100 differential expressed genes and the enrichment of non-zero inflated distributions for the top100 stable genes.

| Platform | Top 100 | Zero-inflated | Not zero-inflated | logOR | 95%CI | p-value |
| --- | --- | --- | --- | --- | --- | --- |
| 10x | Differentially expressed genes | 99 | 1 | 5.19 | 3.36, 8.87 | $3.03 \times 10^{-25}$ |
|  | Stable genes | 35 | 65 |  |  |  |
| Smart-seq | Differentially expressed genes | 97 | 3 | 3.70 | 2.50, 5.36 | $5.46 \times 10^{-18}$ |
|  | Stable genes | 44 | 56 |  |  |  |

### Binarized representation allows for highly scalable analyses

Increasingly larger datasets require increasingly more computational resources. For example, a dataset(29) with 83,262 cells and 20,138 genes after transformation and normalization requires 5.1 Gigabytes of working memory using Seurat(30) and sctransform(31). In contrast, binarizing the same dataset and storing it as bits requires only 300 Megabytes, which is a 17-fold reduction in storage requirements. This potentially boosts scalability of downstream analyses to larger numbers of cells, opening possibilities to get a more fine-grained resolution of biological heterogeneity(32).

## Discussion

In a recent article(8), Jian et al. discuss the “zero-inflation controversy”. They made a distinction between a biological zero, indicating true absence of a transcript, and a non-biological zero, indicating failure of measuring a transcript while it is present in the cell. Likewise, Sakar and Stephens(33) propose to make a distinction between measurement and expression. Following this reasoning, a Poisson model can be used to explain the observed counts, as well as the frequency of observed zero counts. In this case, a non-biological zero encodes useful information, i.e. the gene is unlikely to be highly expressed. However, this model is not suited for the observed zero-inflation. Instead of resuming to a zero-inflated model, we argue that zero-inflation is primarily the result of biological heterogeneity. In other words, the zero-inflation itself is indicative of the multi modal expression of a gene. As such, the binary representation faithfully captures the underlying biology, as the presence of a zero is mainly dictated by the relative abundance of the respective gene.

We showed that analyses based on a binarized representation of scRNAseq data perform on par with count-based analyses. Working with binarizing scRNAseq data has clear additional advantages. The first is simplicity. Binarization circumvents the need to normalize and transform the scRNAseq data. Hence it avoids making various subjective choices and thus improves reproducibility of the subsequent data analyses. Second, binarization alleviate noise, as it is only subject to detection noise opposed to quantification and detection noise(13). Third, binarization reduces the amount of required storage significantly and allows the analysis of significantly larger datasets. This would, for example, allow for a bit implementation of clustering as has been done before in the field of molecular dynamics resulting in a significant reduction of run time and peak memory usage compared to existing methods(34). We have shown that sparsity is inversely correlated with the amount of additional signal that is captured by the quantification of expression. Consequently, binarization will not be useful for all scRNAseq datasets. Previous work suggested that when the detection rate becomes >90% binarization does not perform on par with count-based representation for the task of dimensionality reduction (13). At what detection rate binarizing is not useful anymore for other analyses should be investigated further.

Concluding, as binarized representations of scRNAseq data capture the relevant biological signal, there are opportunities for binarized data analyses methods so that cells can be put into a context of an extremely large number of other cells and be interrogated accordingly.

## Acknowledgements

This research was supported by an NWO Gravitation project: BRAINSCAPES: A Roadmap from Neurogenetics to Neurobiology (NWO: 024.004.012) and the European Union’s Horizon 2020 research.

## Author contributions

GAB, AM, and MJTR conceived the study designed the experiments. GAB performed all experiments and drafted the manuscript. GAB, AM, and MJTR reviewed and approved the manuscript.

## Competing interests

The authors declare no competing interests.

## Notes

### Competing Interest Statement

The authors have declared no competing interest.

